# Nanoparticle-in-Microparticle Oral Delivery System Based on Drug-Loaded Polymeric Micelles

**DOI:** 10.64898/2026.03.17.712272

**Authors:** Hen Moshe Halamish, Roni Sverdlov Arzi, Alejandro Sosnik

## Abstract

This work develops and characterises a hierachichal oral drug delivery system based on the microencapsulation of drug-loaded amphiphilic nanogels within a mucoadhesive alginate/chitosan shell. Results show a more controlled release and a statistically significant oral half-life with respect to the free drug.

---

One of the challenges in pharmaceutical research and development pertains to the poor aqueous solubility and permeability of hydrophobic drugs [1], a property that is common to approximately 50-70% of the drugs on the market [2]. Low solubility in biological fluids leads to limited absorption in the gastrointestinal tract and low oral bioavailability, challenging therapeutic efficacy [3]. Sustained oral drug delivery is difficult to accomplish due to the short residence time and low permeability across the intestinal epithelium of conventional formulations. Polymeric micelles are nanoparticles generated by the self-assembly of polymeric amphiphiles when the concentration in water is above the critical micellar concentration. Depending on the molecular architecture of the amphiphile, they display a hydrophilic corona and a hydrophobic core or more complex nanostructures [4,5] and they have become a valuable nanotechnology platform for the encapsulation, delivery and targeting of small-molecule lipophilic drugs [6,7]. More recently, they have been engineered to fit the encapsulation of biologicals [8]. Historically, these drug nanocarriers have been investigated for intravenous administration though their efficacy in mucosal drug delivery has been proposed [9–11]. We pioneered the use of polymeric micelles in mucosal drug delivery and demonstrated their ability to increase the systemic bioavailability by various administration routes such as oral, intranasal and ocular [12–15]. Despite the advantages shown by oral polymeric micelles, they usually display burst release, and the harsh gastric conditions could degrade the payload before it undergoes intestinal absorption. To prolong the residence time of nano-drug delivery systems in the gastrointestinal tract, we developed Nanoparticle-in-Microparticle Oral Delivery Systems (NiMODS) in which pure drug nanoparticles are microencapsulated within a mucoadhesive matrix [16,17]. These oral delivery systems exhibit improved oral pharmacokinetics characterized by lower plasmatic maximum concentration (C_max_), higher area-under-the-curve (AUC_0-∞_) and longer half-life (t_1/2_) than the free unprocessed and nanonized drugs [15,16]. Aiming to extend the application of this concept, in this work, we investigated a NiMODS comprised of amphiphilic nanogels of a poly(vinyl alcohol)-poly(methyl methacrylate) (PVA-*g*-PMMA) graft copolymer (17% w/w PMMA content) loaded with the tyrosine kinase inhibitor dasatinib (10% w/w payload) and crosslinked with boric acid [18,19], and subsequently encapsulated within alginate matrix microparticles ionotropically crosslinked with calcium cations and chitosan. The drug-loaded nanogels were produced by a one-step process in a Y-shape microfluidics device. The hydrodynamic diameter of freshly prepared dasatinib-loaded nanogels, as measured by dynamic light scattering (DLS), was 149 ± 15 and 63 ± 3 nm by intensity and number, respectively, and the polydispersity index 0.296 ± 0.054 [19]. To encapsulate them within the mucoadhesive microparticles, they were spray-dried utilizing the Büchi Nano Spray Dryer B-90 HP [20] and the dry powder (**Fig. 1A**) was redispersed in a sodium alginate solution immediately before the microencapsulation stage using the Büchi Encapsulator B-390 [17]. The hydrodynamic diameter of spray-dried dasatinib-loaded nanogels upon redispersion in the original volume of water was 129 ± 4 and 54 ± 4 nm by intensity and number, respectively, with a PDI of 0.312 ± 0.004, confirming their very good redispersibility. A decrease in the nanogel size upon spray-drying and redispersion is associated with the consolidation of the crosslinked PVA network upon drying. Finally, a two-stage film-coating of the NiMODS with poly(methacrylate) copolymers conferred gastro-resistance and more selective drug release under the gut conditions. The first film-coating stage comprised Eudragit^®^ RL PO, a water-insoluble copolymer of ethyl acrylate, methyl methacrylate, and a low content of methacrylic acid ester with quaternary ammonium groups that swells becoming water-permeable and designed for sustained drug release. The second film-coating was conducted with Eudragit^®^ S100, a copolymer of methacrylic acid and methyl methacrylate in a 1:2 weight ratio that is insoluble under acidic pH and dissolves at intestinal pH [21]. The rationale for the double film-coating relies on the selective solubility of the outer film in the gut, exposing the inner one that is water-insoluble and permeable and controls the drug release. The uncoated swollen drug-loaded NiMODS were spherical, with smooth surface and a size of ~1.5 μm (**Fig. 1A**). After one and two film-coating cycles, they swelled less and partly lost their spherical shape and surface smoothness (**Fig. 1B,C**). Their diameter, as measured by optical microscopy, was approximately 30-50 μm, with a final dasatinib loading of 1.1% w/w based on dry weight.

**Fig. 1.**
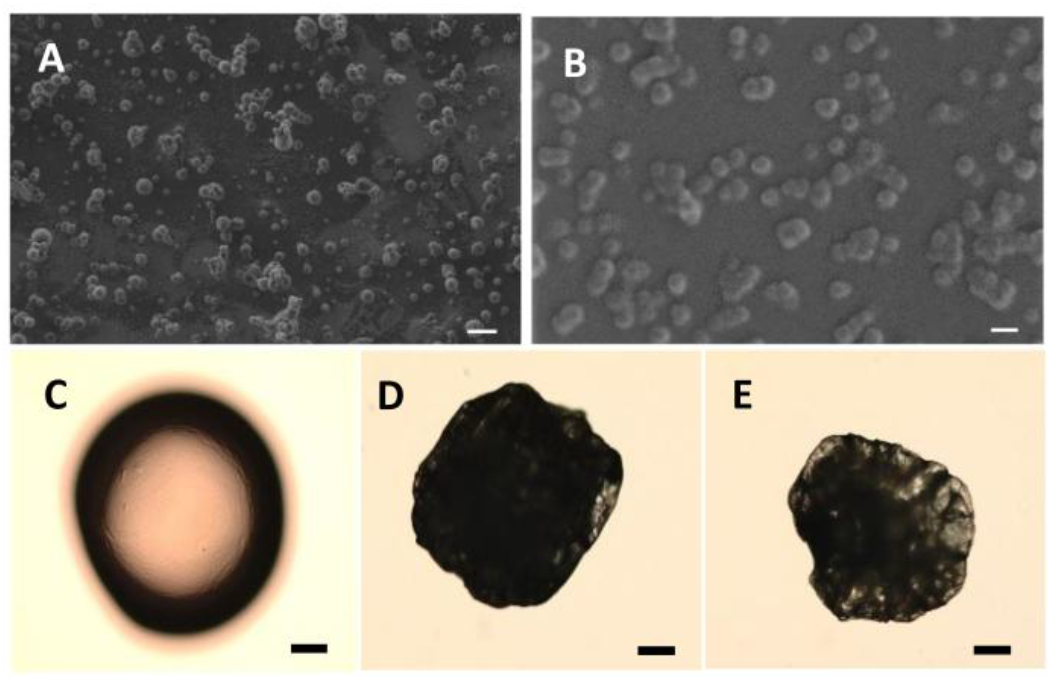
High Resolution-Scanning electron microscopy micrograph of dasatinib-loaded nanogels (A) after spray-drying in the Büchi Nano Spray Dryer B-90 HP and (B) and re-dispersion in water. (C-E) Optical microscopy micrographs of dasatinib-loaded NiMODS produced in the Büchi Encapsulator B-390. (C) Without film-coating, (D) after one film-coating and (E) after two film-coatings. Scale bars: (A) 1 µm, (B) 100 nm and (C-E) 10 µm. The drug loading is 1.1% w/w based on dry weight.

NiMODS are envisioned for oral drug delivery. Thus, two polysaccharides with excellent biocompatibility, namely alginate and chitosan, and classified as “Generally Recognized As Safe” by the US-Food and Drug Administration were used to produce the microparticles [22,23]. However, PVA-*g*-PMMA nanogels could be released in the gut upon disintegration of the microparticulate matrix. The compatibility of the nanogels and their non-crosslinked counterparts was assessed in the Caco-2 and HT29-MTX cell lines for 4-72 h. The former is a human colon carcinoma cells line with an enterocyte-like phenotype, while the latter is a mucin-secreting Goblet cell-like line. These cell lines are a reliable model of the intestinal epithelium *in vitro* [24]. Regardless of the incubation time, cell viability was >78% for both types of nanoparticles, indicating that all the nanogel components including boric acid are cell compatible (**Fig. 2**). These results were with good agreement with previous cell compatibility assays in *vitro* [19,25] and above the value established by the ISO 10993-5 as limit for cell toxicity *in vitro* [26]. The apparent permeability (P_app_) of the 0.1% w/v non-loaded nanogels was evaluated in a model of the intestinal epithelium *in vitro* based on the co-culture of Caco-2 and HT29-MTX cell lines in a 9:1 cell number ratio [27]. The P_app_ was 32.6 ± 3.2 × 10^−7^ cm/s, in good agreement with previous permeability studies *in vitro* with other amphiphilic nanoparticles of similar size [28].

**Fig. 2.**
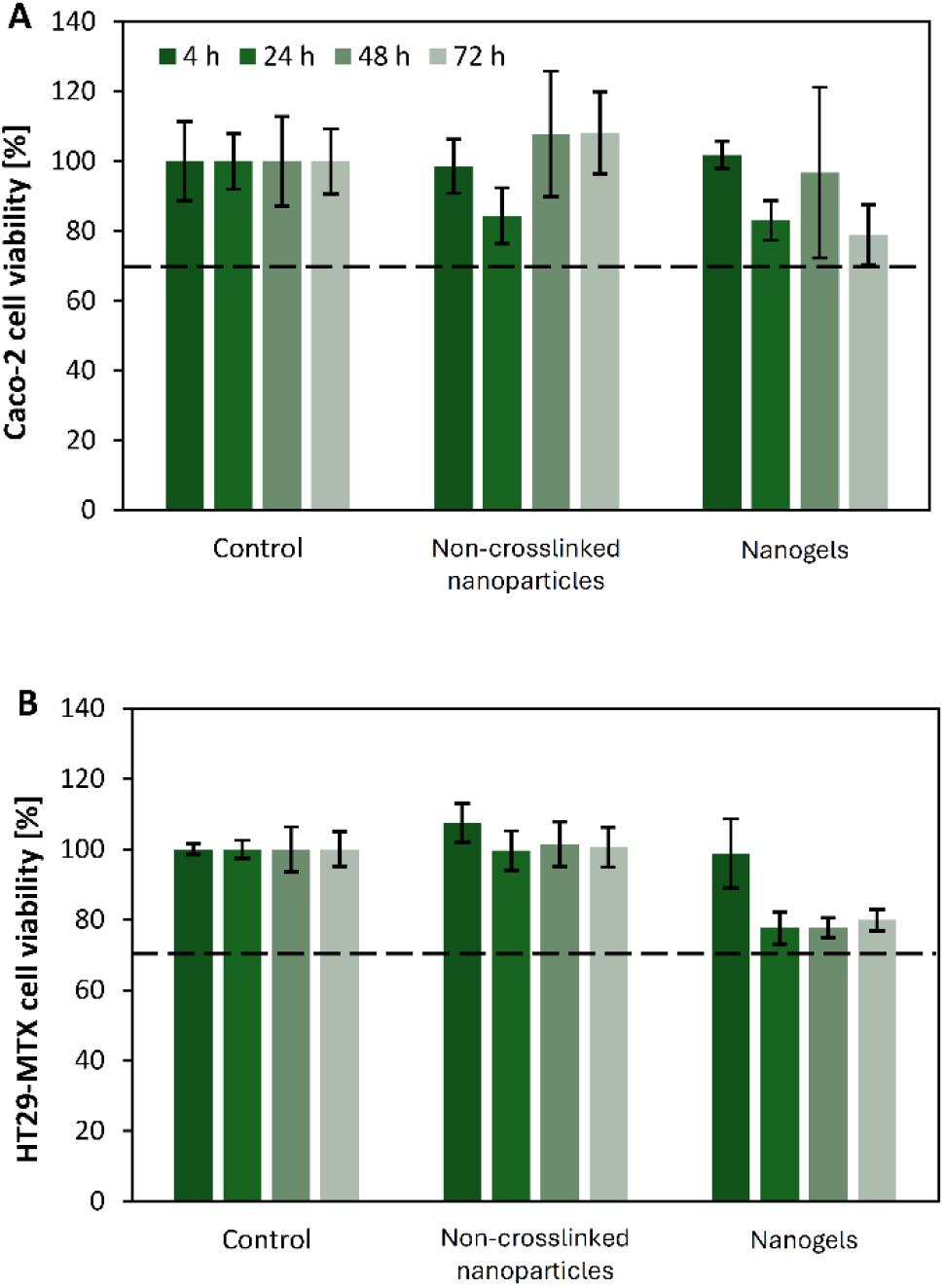
Viability of (A) Caco-2 and (B) HT29-MTX cells after exposure to dasatinib-free non-crosslinked PVA-*g*-PMMA nanoparticles and nanogels for 4, 24, 48 and 72 h at 37°C, as determined by the 3-(4,5-dimethylthiazolyl-2)-2,5-diphenyltetrazolium bromide (MTT) assay (n = 3). The dotted line indicates the 70% cell viability acceptable according to the ISO-10993 guidelines.

The release kinetics of dasatinib from the nanogels and the film-coated NiMODS was assessed under two different pH conditions that mimic the stomach (pH 1.2) and the small intestine (pH 6.8) and compared to that of the free drug. Under both pH conditions, the drug release from PVA-*g*-PMMA nanogels was slower than the release of the free drug (**Fig. S1**); a statistically significant difference was measured only after 4 h for pH 1.2 (p < 0.01) and after 30 min for pH 6.8 (p < 0.05). For example, after 4 h at pH 1.2, the cumulative release of the free drug was 66% compared to 42% of the loaded nanogels (**Fig. S1**). This indicates that these nanocarriers slow down the release kinetics [29]. Interestingly, the release rate of the same formulation in different media did not substantially change. Then, we assessed the dasatinib release from double film-coated NiMODS. Microencapsulation resulted in a more sustained release rate with respect to the free drug and the drug-loaded nanogels at pH 6.8 (**Fig. 3**); 16% of the cargo was released after 4 (p < 0.01). The drug release data were fitted to different release models using the DDSolver add-in program in Microsoft Excel [30], the Weibull model being the best fit to describe cumulative drug release in solution from matrix-type drug delivery systems [31]. These results suggested the sustained release of the cargo upon oral administration and during the gastrointestinal transit.

**Fig. 3.**
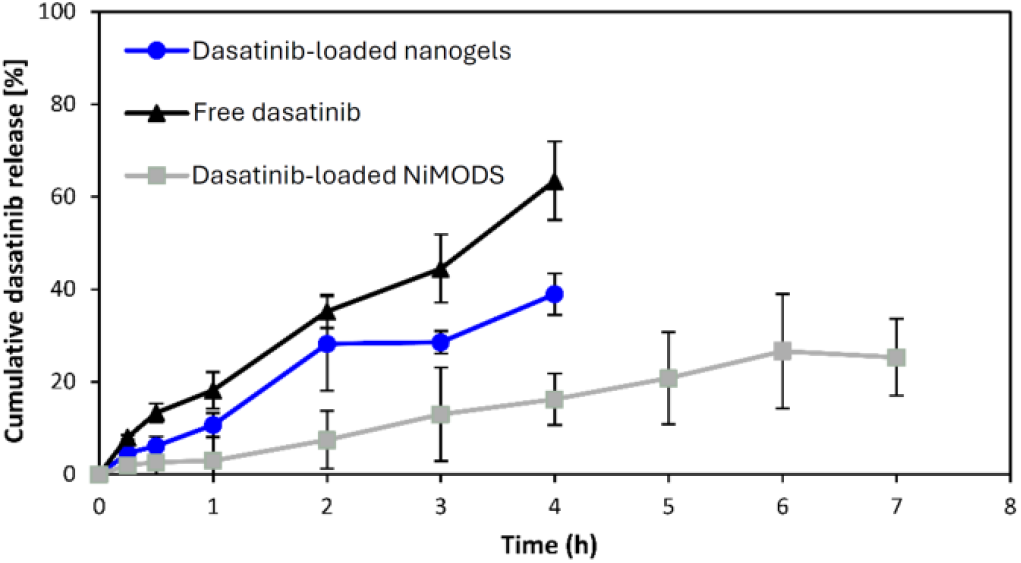
Dasatinib release *in vitro*, at pH 6.8, under sink conditions. NiMODS are double film-coated with poly(methacrylate) copolymers.

The oral pharmacokinetics of the different formulations was assessed in Sprague-Dawley rats (n = 6) and data analysed with PKSolver add-in program [32]. We assessed the performance of unprocessed dasatinib and dasatinib-loaded nanogels at a single dose of 10 mg/kg and of unprocessed dasatinib and double film-coated NiMODS at a single dose of 5 mg/kg.

Administering a dose of 10 mg/kg with dasatinib-loaded NiMODS was not possible in this small animal model due to the relatively low dasatinib loading of this formulation (1.1% w/w) (**Fig. 4**).

**Fig. 4.**
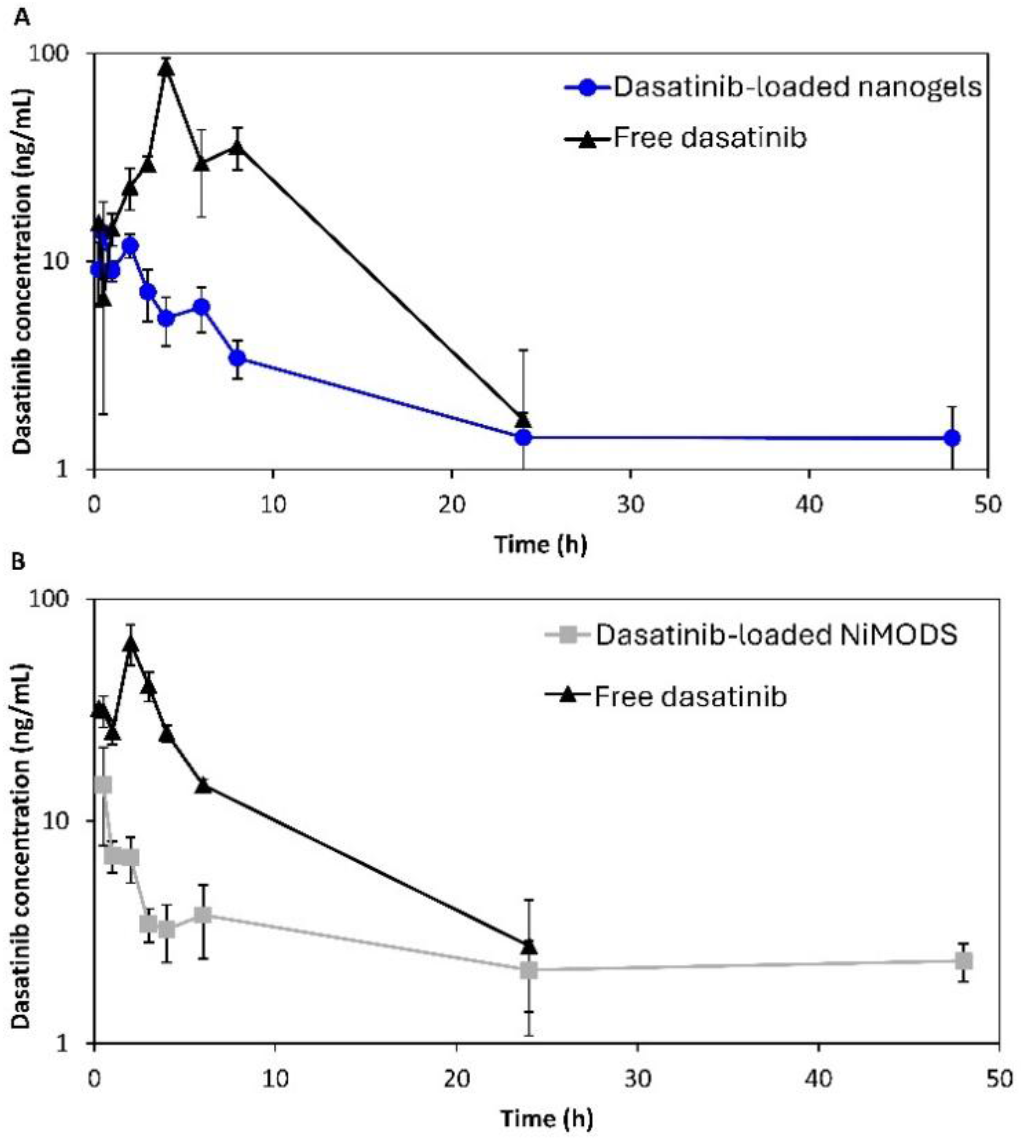
Mean plasma concentration versus time profiles following the oral administration of one single dasatinib dose of (A) 10 mg/kg and (B) 5 mg/kg. Formulations were encapsulated within gelatin capsules that disintegrated in the rat mouth (n = 6).

For a 10 mg/kg dose, unprocessed dasatinib showed a C_max_ of 86.4 ng/mL, a time to the C_max_ (t_max_) of 3.8 h, an area-under-the-curve (AUC_0-∞_) of 604.2 ng/mL/h and a t_1/2_ of 3.8 h (**Fig. 4, Table 1**). For the same dose, dasatinib-loaded nanogels showed a C_max_ of 13.8 ng/mL that was reached 0.5 h post-administration (as opposed to the 4 h showed by the free drug) and the AUC_0-∞_ decreased to 168.9 µg/mL/h, all the differences between the samples being statistically significant (p < 0.01). In addition, the t_1/2_ increased from 3.8 to 19.4 h (**Table 1**). These results indicated that the release of dasatinib from the nanogels takes places at a substantially slower rate than the free drug (in line with the release results *in vitro*), leading to a more moderate and prolonged intestinal absorption. A decrease of the C_max_ could be associated with lower off-target toxicity, though the sharp decline in the AUC_0-∞_ may jeopardize therapeutic efficacy of the loaded nanogels [33]. It is important to point out that dasatinib could be absorbed better in the stomach due to his strong pH-dependent aqueous solubility (18.4 mg/ml at pH 2.6 to 0.008 mg/ml at pH 6.0) [34,35]. Then, we compared the performance of NiMODS with the free drug upon administration of an oral dose of 5 mg/ kg. Owing to the relatively low dasatinib content of the NiMODS (1.1%) a 10 mg/kg dose was unfeasible in rats. As we anticipated, double film-coated NiMDSs showed a statistically significant decrease of C_max_ from 63.3 to 14.6 ng/mL and a dramatic prolongation of the t_1/2_ from 6.7 h to 67.2 h (p < 0.05), suggesting that the microparticles undergo mucoadhesion and control the release kinetics. Furthermore, as opposed to the nanogels, for this dose, the AUC_0-∞_ of both formulations was almost identical (370.0 and 366.3 µg/mL/h for the free and the encapsulated drug, respectively) and statistically insignificant (p > 0.8), as shown in **Table** 1 Remarkably, a 5-mg/kg dose with NiMODS resulted in a 2.2-fold increase of the AUC_0-∞_ with respect to a 10-mg/kg dose with nanogels which strongly suggests the key role played by the mucoadhesive microparticles to ensure a more prolonged residence time of the nanogels in the gut and a more efficient release and absorption with respect to the free drug-loaded nanogels. On one hand, the oral bioavailability, as estimated by the AUC_0-∞_, was substantially lower with the loaded nanogels than with free pure dasatinib nanoparticles [36]. On the other, the C_max_ was more moderate when encapsulated within the microparticles with a similar oral bioavailability which may contribute to lower systemic toxicity [37]. In conclusion, this study demonstrated that the features of both nanoparticles and microparticles can be integrated into hierarchical drug delivery systems for the mucosal drug administration of a broad spectrum of small-molecule and biological drugs, and to tune the release kinetics and improve the bioavailability and therapeutic efficacy [38–41].

**Table 1.** Pharmacokinetic parameters after oral administration of a single dose of unprocessed dasatinib, and dasatinib-loaded nanogels and double film-coated NiMODS to Sprague-Dawley rats (n = 6).

| Pharmacokinetic parameter | Raw dasatinib 10 mg/kg |  | Nanogels 10 mg/kg |  | Raw dasatinib 5 mg/kg |  | NiMODs 5 mg/kg |  |
| --- | --- | --- | --- | --- | --- | --- | --- | --- |
|  | Mean | CV % | Mean | CV % | Mean | CV % | Mean | CV % |
| $C_{max}$ (ng/mL) | 86.4 | 26.7 | 13.8 | 30.2 | 63.3 | 30.4 | 14.6 | 73.9 |
| $AUC_{0-\infty}$ ($\mu\text{g/mL/h}$) | 604.2 | 8.6 | 168.9 | 29.1 | 370.0 | 24.4 | 366.3 | 36.1 |
| $t_{max}$ (h) | 4.0 | 0 | 0.5 | 54.6 | 2.0 | 47.1 | 0.5 | 61.2 |
| $t_{1/2}$ (h) | 3.8 | 45.4 | 19.4 | 21.6 | 6.7 | 63.8 | 67.2 | 46.1 |

## Supporting information

Experimental section

## CRediT authorship contribution statement

Hen Moshe Halamish: Methodology, Investigation, Data curation, Visualization, Validation. Roni Sverdlov Arzi: Methodology, Data curation. Sosnik Alejandro: Original draft, Visualization, Validation, Supervision, Resources, Project administration, Methodology, Investigation, Funding acquisition, Conceptualization.

## Acknowledgements

The authors would like to thank the financial support of the Uzi & Michal Halevy Fund for Innovative Applied Engineering Research for funding the *in vivo* studies conducted in this work. A.S. thanks the support of the Tamara and Harry Handelsman Academic Chair.

## Data availability

The experimental section is included in the supplementary information file.

