## Supplementary material for "Nanoparticle-in-Microparticle Oral Delivery System Based on Drug-Loaded Polymeric Micelles": Experimental section

### Synthesis of poly(vinyl alcohol)-*graft*-poly(methyl methacrylate) copolymer

An amphiphilic PVA-based graft copolymers with poly(methyl methacrylate) (PMMA) as hydrophobic blocks were synthesized by free radical graft polymerization. For this, PVA (0.4 g, Mowiol® 4–88, molecular weight-average of 31,000 g/mol, 87–89% hydrolysis, Sigma-Aldrich, St. Louis, MO, USA) was dissolved in distilled water (100 mL) at room temperature (RT) under gentle magnetic stirring. Tetramethylethylenediamine (0.18 mL, TEMED, Alfa Aesar, Heysham, UK) and nitric acid 70% (0.45 mL, Bio-Lab Ltd., Jerusalem, Israel) were diluted in distilled water (50 mL). The two solutions were degassed in two separate containers using an ultrasonic bath (Elmasonic S30, Elma Schmidbauer GmbH, Singen, Germany) at RT (30 min), mixed and purged with $N_2$ (30 min) at RT under magnetic stirring. At this stage, the original 70% nitric acid (0.45 mL) undergoes a 1:333 dilution (0.45 mL of nitric acid in a total of 150 mL of aqueous medium). This results in a final nitric acid concentration of 0.21%. Then, MMA (99% purity, Alfa Aesar) pre-treated with a basic alumina column (58 Å pore size, Sigma-Aldrich) to remove radical inhibitors before use was dispersed in degassed water (48 mL) and injected into the solution containing PVA, TEMED and nitric acid utilizing a syringe through a septum. The reaction mixture was heated to 35˚C and cerium(IV) ammonium nitrate (0.66 g, CAN, STREM Chemicals, Newburyport, MA, USA) dissolved in degassed water (2 mL) was added. Thus, the final nitric acid concentration during the reaction was 0.1575%. The free radical graft polymerization was allowed to proceed for 2 h at 35ºC and the reaction quenched with hydroquinone (0.182 g, Merck

GmbH, Hohenbrunn, Germany). The crude aqueous solution was dialyzed against water for three days with frequent water exchanges (regenerated cellulose dialysis membranes, molecular weight cut-off of 12–14 kDa, Spectra/Por® 4 Dialysis Membrane; Spectrum Chemical MFG, New Brunswick, NJ, USA), frozen in liquid $N_2$ and freeze-dried for 72 h (Labconco Free Zone 4.5 plus L Benchtop Freeze Dry System, Labconco, Kansas City, MO, USA). A blank reaction (i.e., without the use of the MMA monomer) was carried out as well. The PVA-*g*-PMMA copolymer used in this work contained 17% w/w of PMMA, as quantified by $^{1}$H-Nuclear Magnetic Resonance [1].

For cell permeability studies *in vitro*, PVA-*g*-PMMA was fluorescently labeled by the conjugation of fluorescein isothiocyanate (FITC, Sigma-Aldrich). For this, the copolymer (100 mg) was dissolved in N,N-dimethyl formamide (2 mL, DMF, Bio-Lab Ltd.) under magnetic stirring. Then, FITC was dissolved in DMF (70 mg/mL, 0.2 mL), added to the copolymer solution, and the mixture was stirred (16 h) protected from light, at 32°C. Finally, the product was dialyzed (72 h, regenerated cellulose dialysis membranes, molecular weight cut-off of 3500 Da, Cellular Sup Membrane Filtration Products, Inc., Seguin, TX, USA), freeze-dried (72 h) and stored protected from light at 4°C until use.

**Production of dasatinib-loaded nanogels**

For the nanoencapsulation of dasatinib within PVA-*g*-PMMA nanoparticles, we used a microfluidics device designed and fabricated in our laboratory [2]. The system includes a microfluidics chip and two continuous infusion pumps (Laboratory Syringe Pump, SYP-01, MRC, Kfar Saba, Israel). For this, PVA-*g*-PMMA (0.45% w/v) and dasatinib (0.05% w/v, Carbosynth Ltd., Compton, UK) were dissolved in methanol:water (1.5:0.5 volume ratio) solution; the final dasatinib loading in the nanoparticles being 10% w/w. Then, 2 mL of the copolymer/drug solution was poured into one syringe pump and 18 mL of water into the second one. In the case of crosslinked nanoparticles (nanogels), 5% w/v boric acid (Bio-Lab Ltd.) solution was added to the water. Finally, the copolymer/drug solution and water were injected into the microfluidics chip at flow rates of 0.1 and 0.9 mL/min, respectively, and the nanoparticles collected in a glass vial. To quantify the dasatinib concentration (expressed as % w/v) after the nanoencapsulation and confirm that no drug was lost during the process, an aliquot of the loaded nanoparticles was freeze-dried, re-dissolved in dimethyl sulfoxide (DMSO, Carlo Erba Reagents SAS, Val de Reuil, France) and the absorbance measured in a microplate spectrophotometer (Multiskan™ GO Microplate Spectrophotometer with SkanIt$^{TM}$ software, Thermo Fisher Scientific Oy, Vantaa, Finland) at a wavelength of 322 nm. A calibration curve of dasatinib in DMSO was prepared in the 0.02-0.00009% w/v range and the absorbance of the sample interpolated to calculate the drug loading according to Equation 1

$$\text{Dasatinib concentration}\left[\%\frac{\text{w}}{\text{v}}\right]=\frac{\text{Measured intensity [a.u.]}}{\text{Graph slope}}, \quad (1)$$

The hydrodynamic diameter and the polydispersity index of dasatinib-loaded nanogels were measured immediately after production using dynamic light scattering (DLS, Zetasizer Nano-ZS, Malvern Instruments, Malvern, UK) with 4MW He-Ne laser (λ= 633 nm), digital correlator ZEN3600 and Non-Invasive Back Scatter technology at scattering angle of 173$^{o}$ to the incident beam and data analyzed using CONTIN algorithms (Malvern Instruments).

Spray-drying of dasatinib-loaded nanogels was performed in a Nano Spray Dryer B-90 HP (Büchi Labortechnik AG, Flawil, Switzerland) using an open loop configuration and the following conditions: 80% spraying, 30-32 mbar pressure, 110 kHz frequency, 90% feeding rate and 120-140 L/min airflow rate at a temperature of 105°C. Dry nanogel powders were visualized by High Resolution-Scanning Electron Microscopy (HR-SEM, Zeiss Ultra-Plus SEM, Carl Zeiss NTS GmbH, Oberkochen, Germany). For this, samples were prepared on silicon wafer (cz polished silicon wafers <100> oriented, highly doped N/Arsenic, SHE Europe Ltd., Livingston, UK) on top of carbon tape (Nisshin EM Co. Ltd, Tokyo, Japan). Nanogels were re-dispersed in the original volume of water and reanalyzed by DLS and HR-SEM to confirm their re-dispersibility. Spray-dried nangels were used to prepare dasatinib-loaded NiMODS (see below).

**Cell compatibility of non-crosslinked nanoparticles and nanogels**

Human colon adenocarcinoma cells, Caco-2 (ATCC® HTB-37TM, American Type Culture Collection, Manassas, VA, USA) and HT29-MTX (mucin secreting, originally established at the European Collection of Authenticated Cell Cultures Cat. No. 12040401 and supplied Sigma-Aldrich) cell lines, were cultured in Dulbecco's Modified Eagle's Medium (DMEM, Life Technologies Corp., Carlsbad, CA, USA) supplemented with L-glutamine, 10% heat-inactivated fetal bovine serum (FBS, Sigma-Aldrich) and penicillin/streptomycin (5 mL of a commercial mixture of 100 U/mL penicillin + 100 µg/mL streptomycin per 500 mL medium, Sigma-Aldrich), maintained at 37°C in a humidified 5% $CO_2$ atmosphere and split every 4–5 and 2-3 days Caco-2 and HT29-MTX, respectively. Cells were harvested by trypsinization (trypsin−ethylenediaminetetraacetic acid 0.25% w/v, Sigma-Aldrich) and the number of live cells was quantified by the trypan blue (0.4% w/v, Sigma-Aldrich) exclusion assay. For Caco-2 cell compatibility of non-crosslinked PVA-*g*-PMMA, the cells were cultured in 96-well plates (7.5 × $10^3$ cells per well, in 100 µL) and allowed to attach (96 h). Then, the medium was replaced with 180 µL of fresh medium and 20 µL of 1% PVA-*g*-PMMA solutions in phosphate buffer saline (PBS, pH 7.4) that were sterilized by filtration (0.22 µm Millex-GV, Merck KGaA, Darmstadt, Germany) and incubated at 37°C (12 h) to allow the formation of polymeric NPs to render final concentrations in culture of 0.1% w/v. At different time points (4-72 h), the medium was removed and fresh medium (100 µL) and sterile 3-(4,5-dimethylthiazol-2-yl)-2,5-diphenyl- tetrazolium bromide solution (25 µL, 5 mg/mL, MTT, Sigma- Aldrich) was added. Samples were incubated for 4 h (37°C, 5% $CO_2$), the supernatant was removed, the formazan crystals were dissolved in DMSO (100 µL) and the absorbance was measured at 530 nm with reference at 670 nm using a Microplate Spectrophotometer (Multiskan GO Microplate Spectrophotometer with SkanIt™ software). The percentage of live cells was estimated with respect to a control treated only with culture medium that was considered 100% viable. The assay for crosslinked counterparts was similar except for a slight difference in the preparation of the samples. In this case, each nanogel suspension was prepared ahead of time with 180 µL medium and 20 µL of sterile copolymer solution (the same final concentration as for non-crosslinked polymeric nanoparticles) and were incubated at 37°C for 12 h to allow their formation. Then, 1 µL of boric acid (5% w/v) was added to the suspension and incubated at 37°C for an additional 12 h to ensure crosslinking. Finally, the culture medium was replaced by the polymeric NPs suspension and the compatibility measured as described above. HT29-MTX cell line compatibility with PVA-*g*-PMMA copolymers was conducted the same way as Caco-2 cell line, except for the number of cultured cells per well (3.5 × $10^3$ cells).

**Nanogel permeability *in vitro***

A preliminary permeability assay was conducted with dasatinib-free PVA-*g*-PMMA nanogels in a model of intestinal epithelium. Transport experiments were performed 10–25 days post-seeding of Caco-2:HT29-MTX co-culture (9:1 cell number ratio) in cell culture inserts (ThinCert™, culture surface of 113.1 $mm^2$, 3.0 µm pore size, Greiner Bio-One GmbH, Frickenhausen, Germany) maintained in 12-well plates (15.85 mm diameter, 16.25 mm height, Greiner CELLSTAR, Monroe, NC, USA) with 0.5 and 1.5 mL of DMEM medium (see above) in the apical and basolateral compartment, respectively. The total number of cells was always $3 \times 10^5$ cells per well. The culture medium was replaced every 2–3 days and the integrity of the cell monolayer was characterized by transepithelial electrical resistance (TEER) measurements performed with an epithelial volt-ohmmeter (“$EVOM^2$”, WPI, Sarasota, FL, USA) (Figure 3-4). For these experiments, only inserts where the resistance was higher than 200 $\Omega \cdot cm^{-2}$ were used. Fluorescein-labeled nanogels were prepared in Hank’s Balanced Salt Solution (HBSS, Sigma-Aldrich) buffer as described above by combining a weight ratio of unlabeled:labeled copolymer of 8:2 and rendered a final concentration of 0.1% w/v. Initially, the medium in the apical (0.5 mL) and basolateral (1.5 mL) was replaced with transport medium (HBSS) and cells were incubated for 15 min at 37°C in a humidified 5% $CO_2$ atmosphere. Then, transport medium in the donor (apical) compartment was replaced by the corresponding sample (0.4 mL) and in the acceptor compartment (basolateral) by fresh transport medium (1.2 mL). After 5, 10, 15, 30, 45, 60, 90, 120, 180, and 240 min, 600 µL of medium was extracted from the basolateral compartment and replaced by the same volume of fresh transport medium maintaining the total volume in the well constant. The quantification of the transported nanogels in the extracted medium was measured by fluorescence spectrophotometry (Fluoroskan Ascent Plate Reader, Thermo Fisher Scientific Oy) using black 96-well flat bottom plates (Greiner Bio-One, Kremsmünster, Austria) at wavelengths of 355 nm for excitation and 538 nm for emission. The measured fluorescence was interpolated according to a calibration curve of the copolymer in HBSS with concentration between 0.0001% and 0.1% w/v ($R^2 = 0.9995$). After the last point (240 min), 50 µL was also removed from the apical side of each sample to calculate the mass balance. The apparent permeability ($P_{app}$) was calculated according to Equation 2

$$P_{app} = \frac{dc}{dt} \cdot \frac{1}{A \cdot C_0} \left[\frac{cm}{s}\right] \qquad (2)$$

Where $\frac{dc}{dt}$ is the permeability rate (µg/s) across the monolayer, $C_0$ is the initial concentration in the donor compartment (µg/$cm^3$), and A is the surface area of the membrane $cm^2$. Results are expressed as mean ± S.D. of at least three experiments.

**Preparation of dasatinib-loaded NiMODS**

Dasatinib-loaded NiMDOS were produced using the Encapsulator B-390 (Büchi Labortechnik AG) which utilizes prilling by vibration technology to form the droplets. For this, low viscosity alginate sodium salt (0.8 g, Alfa Aesar) and ultra-low molecular weight chitosan (0.2 g, molecular weight of 20,000 g/mol, Glentham Life Sciences, UK) were solubilized separately in distilled water (100 mL) under a magnetic stirring. Nitric acid 70% (0.225 mL) was added to the chitosan solution to reach full solubilization and the two solutions were left under magnetic stirring overnight. Calcium chloride (1.11 g, J.T.Baker, Avantor Performance Materials, Gliwice, Poland) was added to the chitosan solution one hour before the crosslinking. Spray-dried dasatinib PVA-*g*-PMMA nanogels (40 mg) were dispersed in the alginate solution (40 mL), obtaining a final drug concentration in the system of 0.1% w/v. The nanosuspension in alginate was fed into the encapsulator to produce the microparticles. A nozzle with an inner diameter of 450 μm diameter was used. The air pressure set for the encapsulation was 1.5 bar, the system pressure was adjusted between 0-300 bar, frequency of vibrations and the voltage applied were adjusted slightly until spherical particles were formed. Then, the microparticles were ionotropically crosslinked in the chitosan/calcium chloride solution and left under gentle magnetic stirring for 30 min. To separate the microparticles from the chitosan solution and to remove drug and calcium chloride residues, they were rinsed with distilled water and filtered through a 150 MM Whatman™ filtration paper (Merck KGaA, Darmstadt, Germany).

Alginate microparticles were suspended in distilled water (40 mL) and immediately frozen in liquid $N_2$ to preserve their shape and size through the drying process and to prevent the drug release from the particles. Then, the microparticles were freeze-dried (Freeze Dryer Unit Beta 1-8 LDplus, Christ, Osterode, Germany) for 72 h. The particles were film-coated with copolymers of the Eudragit® family (Evonik Rohm GmbH, Kirschenallee, Germany). First, both Eudragit® copolymers were dissolved in ethanol under magnetic stirring overnight. Then, the dry microparticles were placed in pure water under magnetic stirring (100 RPM, 3 min) to enable partial swelling and the water excess was removed by a pipette. An Eudragit® RL PO solution (5% w/v, a copolymer of ethyl acrylate, methyl methacrylate, and a low content of methacrylic acid ester with quaternary ammonium groups) was poured on the microparticles and stirred (100 RPM) for 10 min. The excess solution was removed by a pipette and the microparticles were dried for 2 h at RT. Finally, microparticles were immersed in an Eudragit® S100 solution (5% w/v, a copolymer of methacrylic acid and methyl methacrylate in a 1:2 ratio and magnetically stirred at 100 RPM for 10 min and dried overnight.

Dasatinib-loaded NiMODS without film-coating, and Eudragit® RLPO film-coating or Eudragit® RL PO- Eudragit® S100 double film-coating, were visualized under an optical microscope (BX51 Light Microscope, Olympus, Waltham, MA, USA).

For the evaluation of the amount of drug in the microparticles, the percentage of drug residues in the crosslinking solution was measured. For this, 40 mL of the crosslinking solution was freeze-dried, dissolved in dimethyl sulfoxide (5 mL, DMSO, Carlo Erba Reagents, Emmendingen, Germany) and the drug concentration was measured at 309 nm and 322 nm wavelengths using a Multiskan GO Microplate Spectrophotometer with SkanIt™ software (Thermo Fisher Scientific Oy, Vantaa, Finland).

### Dasatinib release from nanogels and NiMODS *in vitro*

The release kinetics of dasatinib from the nanogels and NiMODS under simulated physiological conditions of the gastrointestinal tract (both gastric pH of 1.2 and intestinal a pH of 6.8, 37°C) was assessed by the dialysis membrane method using a dissolution bath (708-DS, Agilent Technologies, Santa Clara, CA, USA) under sink conditions and compared to that of the free drug. Release data were fitted to different mathematical models using DDSolver, an add-in program in Microsoft Excel [3]. For the preparation of a release medium of 6.8, we mixed 50.3 mL of $KH_2PO_4$ solution (68.045 g of $KH_2PO_4$ salt dissolved in 500 mL of water, Merck GmbH, Darmstadt, Germany) and 49.7 mL of $K_2HPO_4$ solution (87.09 g of $K_2HPO_4$ salt dissolved in 500 mL of water, Spectrum chemical MFG Corp., Gardena, CA, USA). The acidic medium of pH 1.2 was a dilution of HCl solution in water (hydrochloric acid 32%, 10M diluted to 0.063M pH 1.2).

First, a calibration curve of dasatinib in the 0.02-0.00001% w/v concentration range was prepared in each medium (gastric- and intestinal-like) and the absorbance of the sample interpolated to calculate the drug loading according to Equation 1. For this, a dasatinib stock solution was prepared in methanol (1 mg/mL) and each of the mediums were prepared as described above. For each release study, we used 0.3 mg of dasatinib in 50 mL of release medium and a total volume of release medium of 850 mL to maintain sink conditions. The dry nanogels and microparticles were dispersed in 50 mL of the pre-heated medium and additionally pure drug was also dissolved in 50 mL of this medium for comparison. These dispersions were poured into dialysis bags (Spectra/Por, molecular weight cut-off of 12-14 kDa) and immersed into release medium baths (850 mL) at 37 ± 1°C under a mechanical stirring (50 RPM) for 6 h (n = 3). At predetermined time points, aliquots (10 mL) of the release medium were withdrawn and replaced by fresh pre-heated medium. This drug/release medium ratio together with the medium exchanges along the experiment ensured the maintenance of sink conditions that are critical in release studies of poorly water-soluble drugs in vitro. Each aliquot (200 µL) was measured in a Multiskan GO Microplate Spectrophotometer with SkanIt™ software at 309 nm and 322 nm wave lengths to determine the amount of released drug.

**Oral pharmacokinetics of dasatinib**

The oral pharmacokinetics of dasatinib was assessed in male Sprague-Dawley (Envigo) following the Directive 2010/63/EU on the protection of animals used for scientific purposes (22 September 2010, European Union) and approval of preclinical protocols by the Committee of Animal Welfare of Technion (IL-035-03-18 and IL-013-01-22). Three dasatinib formulations were tested: (i) free drug, (ii) drug-loaded nanogels and (iii) drug-loaded NiMODS. We used a new administration technique that was developed in our lab [4]. For this, twenty-four animals (weight of 200-280 g) were divided into four groups (n = 6) and trained to swallow empty gelatin capsules. Then, formulations containing the exact drug dose based on the individual weight of each rat (10 mg/kg or 5 mg/kg of dasatinib) were weighed and loaded into 9el gelatin capsules (diameter of 2.7 mm and length of 23 mm, capsule body capacity of 80 µL, Torpac, Inc., Fairfield, NJ, USA) shortly before administration. Next, the capsule was administered to the cheek of the rat using a metallic plunger (Torpac®, Inc.), minimizing animal discomfort. Then, animals received 1 mL of water that dissolved the capsule in the mouth and enabled the swallowing of the formulation. Animals were divided into two groups (n = 3) to allow them social interaction ensuring low stress environment. Blood samples were taken at predetermined time points, immediately centrifuged (3000 RPM, 4ºC for 2 min) to separate the plasma and froze at -80ºC until extraction and analysis by high-performance liquid chromatography (HPLC). The quantification of dasatinib in plasma was carried out by reversed phase HPLC in an Alliance separation module e2695 (Waters Corp. Milford, MA, USA) equipped with an XBridge C18column (3.5 µm, 4.6x250 mm, Waters Corp., Dublin, Ireland) and UV-Vis detector (2998 Photoiodide Array UV/Vis 2D detector, W2998, Waters Corp.). The conditions for the HPLC run were optimized to a final condition of 0.6 mL/min isocratic flow at 25ºC. Each sample was transferred into HPLC vial inserts with bottom spring (300 µL, 6 X 29 mm,

Waters Crop.) and 20 µL were collected from each sample and were injected into the HPLC. Plasma analysis mobile phase consisted of 0.02 M buffer (pH 7.4) and HPLC grade methanol in a volume ratio of 20:80 respectively. The analysis was according to dasatinib calibration curve in plasma in concentration range of 12.5-400 ng/mL ($R^2 > 0.98$). For this, standard stock solution of dasatinib was prepared in HPLC grade menthol to yield concentration of 1 mg/mL and diluted to 4000 ng/mL working solution. Next, 20 µL working solution was spiked with 180 µL plasma to yield dasatinib concentration of 400 ng/mL and then 100 µL were diluted with 100 µL plasma and so on until reaching final concentration of 12.5 ng/mL. For oral pharmacokinetic analysis, 30 µL plasma samples were transferred into polypropylene microcentrifuge tube and mixed with 100 µL cold acetonitrile for protein precipitation, vortexed for 1 min and centrifuged at 12,000 rpm for 12 min at 4ºC (HERMLE Z326K, HERMLE Labortechnik GmbH, Germany). Finally, 100 µL of the supernatant was transferred into HPLC vial and loaded into autosampler tray. All plasma samples preparation was conducted under cool environment. Pharmacokinetic parameters calculations were performed by non-compartmental analysis using PKSolver, an add-in program in Microsoft Excel [5]. The following parameters were calculated, maximum plasma concentration that was defined as $C_{max}$ with $t_{max}$ being the time at which $C_{max}$ was reached, the area-under-the-curve ($AUC_{0-\infty}$) plasma concentration-time between 0 and ∞ and apparent half-life in plasma ($t_{1/2}$).

**Statistical Analysis**

Data were analyzed by relevant statistical tests (two-sided T-test) to determine the difference among groups and treatments. The value of $p < 0.05$ was considered statistically significant. One asterisk (*) indicates p-value smaller than 0.05, two asterisks (**) indicate p-value smaller than 0.01 ($p < 0.01$) and three asterisks (***) indicate p-value smaller than 0.001 ($p < 0.001$).

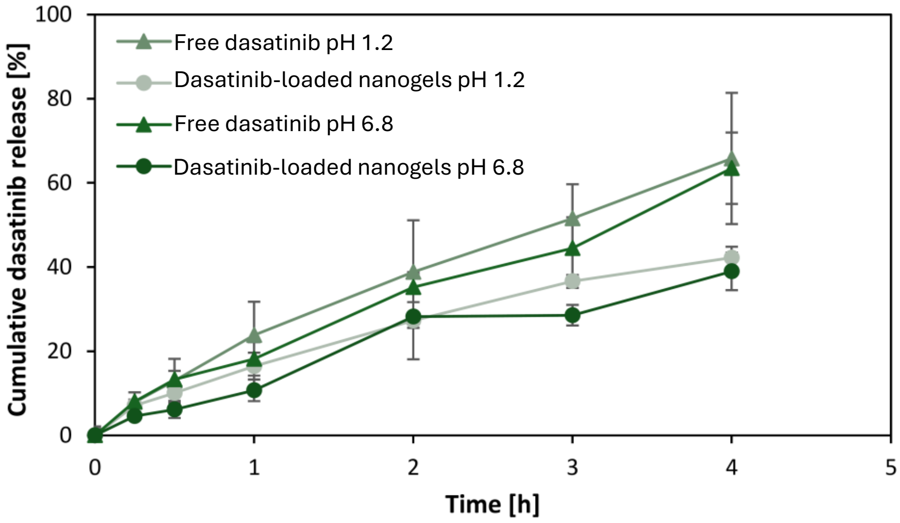


**Fig. S1.** Dasatinib release *in vitro*, at pH 1.2 and 6.8, under sink conditions.
